# Developmental phenomics suggests that H3K4 monomethylation catalyzed by Trr functions as a phenotypic capacitor

**DOI:** 10.1101/2022.03.15.484407

**Authors:** Lautaro Gandara, Albert Tsai, Mans Ekelöf, Rafael Galupa, Ella Preger-Ben Noon, Theodore Alexandrov, Justin Crocker

## Abstract

How epigenetic modulators of gene regulation affect the development and evolution of animals has been difficult to ascertain. Despite the widespread presence of histone 3 lysine 4 monomethylation (H3K4me1) on enhancers, hypomethylation appears to have minor effects on animal development and viability. In this study, we performed quantitative, unbiased and multi-dimensional explorations of key phenotypes on *Drosophila melanogaster* with genetically induced hypomethylation. Hypomethylation reduced transcription factor enrichment in nuclear microenvironments, leading to reduced gene expression, and phenotypes outside of standard laboratory conditions. Our *developmental phenomics* survey further showed that H3K4me1 hypomethylation led to context-dependent changes in morphology, metabolism, and behavior. Therefore, H3K4me1 may contribute to phenotypic evolution as a phenotypic capacitor by buffering the effects of chance, genotypes and environmental conditions on transcriptional enhancers.

**Quote:** *“Developmental biologists are often not so much opposed to a role for ecology as they simply ignore it” –Doug Erwin*^1^

## Introduction

Gene regulation across animal development is carried out by networks of interacting transcription factors, modified by the cellular environment, biochemical pathways, metabolic state, and epigenetic landscapes^2^. These complex regulatory networks are the products of evolution, subject to continual change in response to variable ecologies^3^. To gain traction into this complexity, a classical approach in developmental biology has been to simplify the system using lab-bred model organisms, standardize the experiments under controlled laboratory conditions, and measure pre-defined variables that are expected to change. Such experiments have provided a wealth of information focused on dissecting essential components and their interactions across development.

While research programs with a reductionist focus are powerful, such approaches may not give us the full picture of how systems function in their native environments^4^. Recent advances in high-throughput phenotyping technologies have facilitated a complementary approach: the unbiased and unconstrained exploration of several phenotypes and environmental conditions. New methods in mass spectrometry or automated video tracking are enabling quantitative and cost-effective explorations of complex phenotypes such as metabolism and behavior^5^. Furthermore, acquisition of high-dimensional phenotypic data, or “Phenomics”^5^, could study nuanced modulators of gene expression or robustness-conferring elements, revealing their impacts at the scale of the entire organism and populations.

The monomethylation of histone H3 on lysine 4 (H3K4me1) is an epigenetic mark with disputed roles^6^ in gene regulation—while it has been associated with enhancer elements^7–9^ across the genome of different species^9–11^, its loss appears to have minor effects. In mouse embryonic stem cells, the loss of H3K4me1 from enhancers in Mll3/4 catalytically deficient cells led to minimal effects on transcription^12,13^ and did not disrupt self-renewal^12^. Whilst recent works suggest that H3K4me1 could be relevant in mouse development^14,15^, H3K4me1 hypomethylation produced by disrupting the catalytic activity of Trithorax-related (Trr), the main methyltransferase behind this epigenetic mark in *Drosophila melanogaster*^16^, did not affect development or viability in this species^17^. The lack of clearly defective phenotypes under standard laboratory conditions has therefore led to the hypothesis that H3K4me1 tunes enhancers for a more nuanced response to environmental or genetic stresses^17,18^. However, this subtle effect contrasts with the presence of the epigenetic mark throughout the *Drosophila* genome^6,9^.

To explore comprehensively the effect of H3K4me1 on phenotypes, we designed a developmental phenomics^5^ workflow, and applied it on a H3K4me1 hypomethylation *D. melanogaster* line challenged by various genetic and environmental conditions. Starting from a single regulatory network in hypomethylated embryos, we demonstrated that H3K4me1 may confer transcriptional robustness by localizing transcription factors in nuclear microenvironments. Then we explored the impact of H3K4me1 hypomethylation on a biological system by performing multiple phenotypic assays on larvae on which a catalytically impaired version of Trr led to H3K4me1 hypomethylation. Consistent with the ubiquitous presence of H3K4me1 across the genome^8,9,19^, hypomethylation triggered changes in morphology, metabolism, behavior, and adaptability in response to genetic and environmental challenges. In sum, global H3K4me1 hypomethylation led to reduced developmental robustness and revealed phenotypic variation depending on environmental and genetic contexts, supporting the hypothesis that H3K4me1 acts as a phenotypic capacitor.

### Transcriptionally active *shavenbaby* (*svb*) loci have locally enriched levels of H3K4me1

Previous works showed that global patterns of gene expression were unaffected by H3K4me1 hypomethylation^12,17^. However, H3K4me1 exhibits clearly different trends between its global nuclear distribution and enrichment around individual genes compared to other histone modifications during embryo development^20^, potentially suggesting that it serves specific regulatory functions at those locations. As an increase in H3K4me1 is associated with active enhancers^9^ and with the activity of most members in the *svb* network (Supplementary Figure 1A, see *Correlation between H3K4me1 deposition and the regulation of the svb network* in **Materials and Methods**), we investigated if transcriptionally active *svb* enhancers show enrichment for H3K4me1. This extensively-studied regulatory network controls the differentiation of ectodermal cells into trichomes in late embryonic stages. To capture cells where *svb*-related ventral enhancers are active, we FACS-sorted nuclei from stage 15 *Drosophila melanogaster* embryos from a line with reporter genes driven by different *svb* enhancers. The *“E10” svb* enhancer drives the expression of GFP, whilst *dsRed* is driven by the “7” enhancer (Figure 1A). The sorted nuclei were then processed through ChIP-Seq targeting H3K4me1 (Figure 1B and Supplementary Figure 1B). As expected, H3K4me1 marked all the known embryonic enhancers of *svb* in nuclei from the entire embryo (“All”, Figure 1B). Cells where the reporter gene for a specific *svb* enhancer is active (“7” or “E10”, Figure 1B) showed slightly increased levels of monomethylation over the corresponding enhancer and across the *svb* regulatory region.

**Figure 1.**
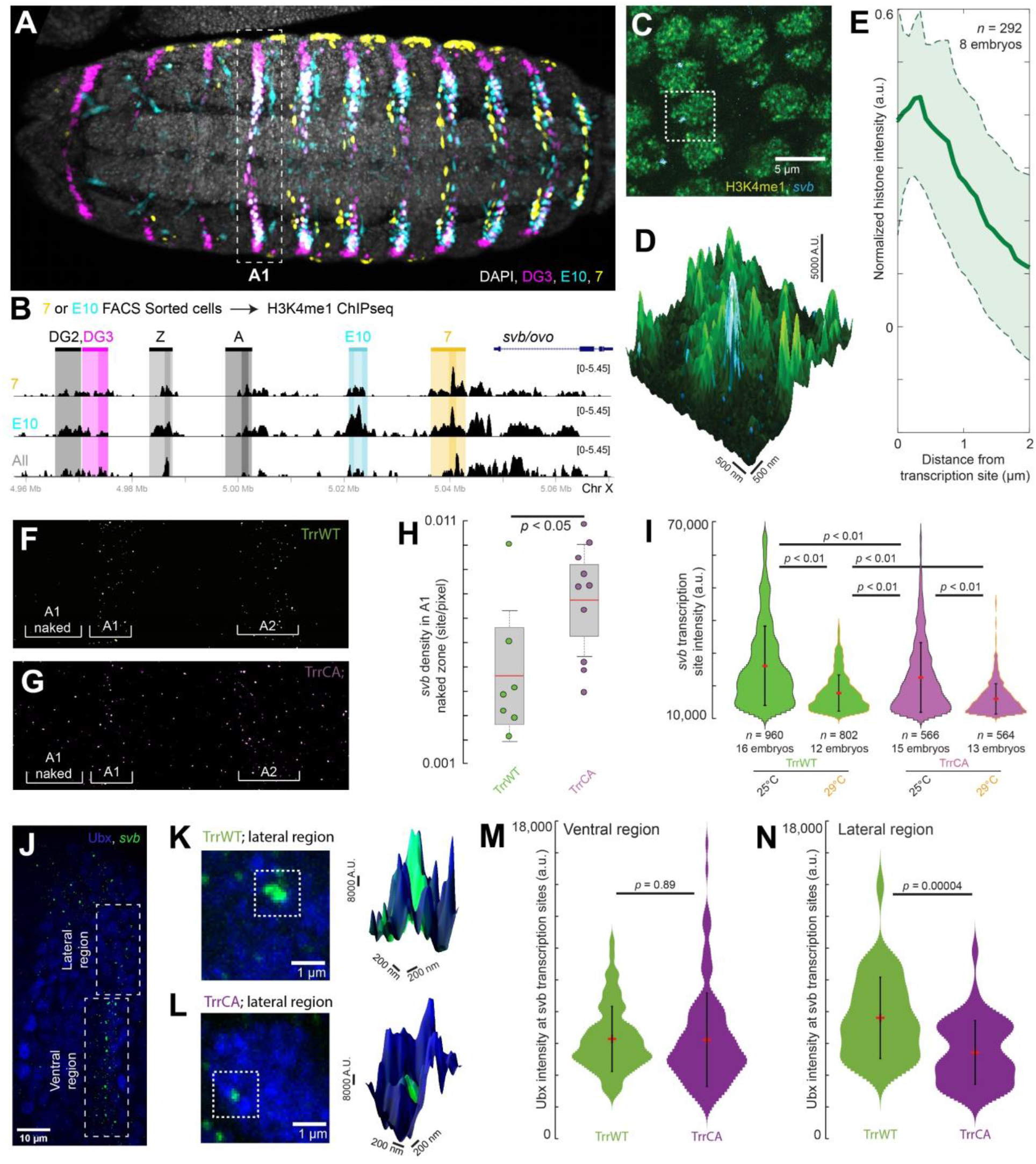
H3K4me1 at the svb locus improves the robustness of transcriptional output through the regulation of transcriptional microenvironments. Related to Supplementary Figure 1. (A) Ventral view of a stage 15 *Drosophila melanogaster* embryo from the fly line used for the ChIP-Seq experiment stained for the products of the reporter genes driven by the *svb* enhancers *DG3, E10*, and *7*. The white dotted box shows the location of the A1 segment along the embryo. (B) ChIP-Seq experiment using cells from stage 15 *Drosophila melanogaster* embryos sorted by reporter gene activity. The panel shows H3K4me1 enrichment on a portion of the *svb/ovo* locus, on cells with an active *7* enhancer (orange), an active *E10* enhancer (cyan), or cells from the entire embryo (“All”, gray). (C) High resolution confocal imaging experiments in stage 15 *w*^*1118*^ embryos show that active *svb* transcription sites are in regions with enriched levels of H3K4me1. (D) Zoomed-in view of the dotted box in (C) with the height indicating the intensity of the H3K4me1 signal. (E) Normalized average H1K4me1 intensity over 292 transcription sites in 8 embryos. The shaded region is one standard deviation (s.d.). (F & G) *svb* transcription sites at 25 °C on the ventral side of the first two abdominal (A1 & A2) segments of stage 15 embryos in both *trr*^*1*^ lines. (H) Transcription site density in front of the A1 ventral band (“A1 naked”). Number of embryos: 7 (TrrWT) and 10 (TrrCA). The boxed region is one s.d. and the tails are two s.d. (95 %). (I) Intensity of *svb* transcription sites at different temperatures. The red dot is the mean and the bar is one s.d. (J) Confocal microscopy image of active *svb* transcription sites and Ubx distribution in the first abdominal segment of a stage 15 TrrWT embryo. (K & L) High resolution confocal imaging experiments in stage 15 embryos show that H3K4 hypomethylation impairs Ubx recruitment to svb transcription sites. Right panels: Zoomed-in view of the dotted boxes with the height indicating the intensity of the Ubx signal. (M & N) Intensity of the Ubx signal in *svb* transcription sites, measured exclusively in the ventral (M) or in the DG3-only lateral region (N). The red dot is the mean and the bar is one s.d. Number of embryos: TrrWT = 5 embryos, TrrCA = 8 embryos. Number of analyzed transcription sites in the ventral region (M): TrrWT n = 69, TrrCA = 139, and in the lateral region (N): TrrWT n = 38, TrrCA n = 45.

We performed high-resolution confocal imaging along the ectoderm in the first abdominal (A1) segment (white box in Figure 1A) of stage 15 embryos (*w*^*1118*^) to see if H3K4me1 is locally enriched at active *svb* loci. We located cells that are expressing *svb* using fluorescence *in situ* hybridization (FISH) with RNA probes targeting the *svb* mRNA^21^ and stained for H3K4me1 using immunofluorescence (IF) (Figure 1C & D, *Sample preparation and staining for confocal imaging* in **Materials and Methods**). Plotting the average radial intensity of H3K4me1 as a function of distance from the transcription site showed that *svb* transcription sites sit on or near a local maximum (Figure 1E), reminiscent of localized Ubx concentrations around the same locus^22^. This local enrichment of the mark was stronger than previously observed at *hb* transcription sites in a previous work^20^ (Supplementary Figure 1C & D, adapted from Tsai & Crocker, 2022). Thus, the enrichment we observed with both ChIP-Seq and imaging suggests that H3K4me1 is locally enriched at transcriptionally active *svb* enhancers.

### Hypo-monomethylation of H3K4 lowered the transcriptional output of *svb*

To identify the effects of losing H3K4me1 on *svb* expression, we disrupted the catalytic activity of Trr. We used a previously characterized fly line with the *trr*^*1*^ null allele, complemented with a construct bearing a cysteine-to-alanine (C2398A) mutation ((*trr*^*1*^*;;trr*(C2398A)), “TrrCA”), which led to a specific reduction in H3K4me1 deposition, but rescued the *trr*^*1*^-induced lethality^17^. This TrrCA line produced fertile adults with a normal life span, no gross morphological abnormalities, and normal gene expression in adult brains and larval wing imaginal discs compared to control lines^17^. We used the *trr*^*1*^ null line rescued with the wild-type Trr ((*trr*^*1*^*;;trr*(WT)), “TrrWT”) as our control to rule out effects from the *trr*^*1*^ line itself.

To observe how hypomethylation changes *svb* regulation, we quantified *svb* transcription sites in the A1 segment of stage 15 embryos using FISH. Even at room temperature (25 °C), the TrrCA line had numerous transcription sites outside of the ventral stripes, while there were few in the TrrWT line (Figure 1F & G): the region in front of the A1 stripe had an average of 0.0077 sites per pixel in the hypomethylation line, versus 0.0046 in the wild-type (*p* < 0.05 two-tailed Student’s *t*-test, Figure 1H). While the densities of transcription sites within the A1 ventral stripe were similar between TrrCA and TrrWT (Supplementary Figure 1E), the intensity of *svb* transcription sites in A1 of TrrCA was lower than TrrWT (*p* < 0.01) at 25 °C (Figure 1I). At 29 °C, *svb* transcription site intensity decreased for both TrrCA and TrrWT; however, it was again lower in the TrrCA line (*p* < 0.01) (Figure 1I).

These *svb* transcription sites have previously been characterized as being inside of transcriptional microenvironments, which are locally enriched for TFs required for *svb* expression^22^. Thus, we analyzed if hypomethylation affects the local enrichment of the transcription factor Ubx, the Hox factor driving ventral *svb* expression in the A1 segment (Figure 1J-N). Despite the difference in *svb* intensity, Ubx intensity at *svb* transcription sites was not different between the TrrWT and TrrCA lines in the ventral region of the *svb* expression pattern (Figure 1J & M). However, in the lateral region, where *svb* transcription is driven by only a single enhancer, *DG3*, and trichome development is less robust^21^, Ubx intensity was significantly reduced by H3K4me1 hypomethylation (Figure 1 K, L & N). In sum, H3K4me1 hypomethylation reduced both the accuracy and levels of *svb* expression and, in the absence of enhancer redundancy, impaired local *svb* transcriptional environments.

### Reduced H3K4me1 impaired the robustness of trichome phenotype at increased temperatures

Nuclear microenvironments are essential for preserving transcriptional activity from the effects of both environmental and genetic perturbations^21,23^. Based on the effect of H3K4me1 hypomethylation on *svb* transcriptional microenvironments, we analyzed the robustness of trichome development.

At room temperature (25 °C), TrrWT larvae had no trichomes outside of the ventral band (Figure 2A). In contrast, TrrCA larvae developed extra trichomes outside of the normal ventral band (Figure 2B, arrows). While TrrWT larvae had similar numbers of A1 ventral trichomes at 25 °C, 29 °C, and 32 °C, the number of trichomes progressively dropped in the TrrCA line as the temperature increased (p < 0.05, two-tailed Student’s *t*-test) (Figure 2C). Similar trends were observed at the lateral edge; however, the TrrCA line had fewer trichomes than TrrWT even at room temperature (Figure 2D). These results indicate that H3K4me1 hypomethylation reduces the robustness of the trichome pattern at increased temperatures, which is consistent with previous observations showing that H3K4 hypomethylation leads to environment-dependent phenotype alterations^17^.

**Figure 2.**
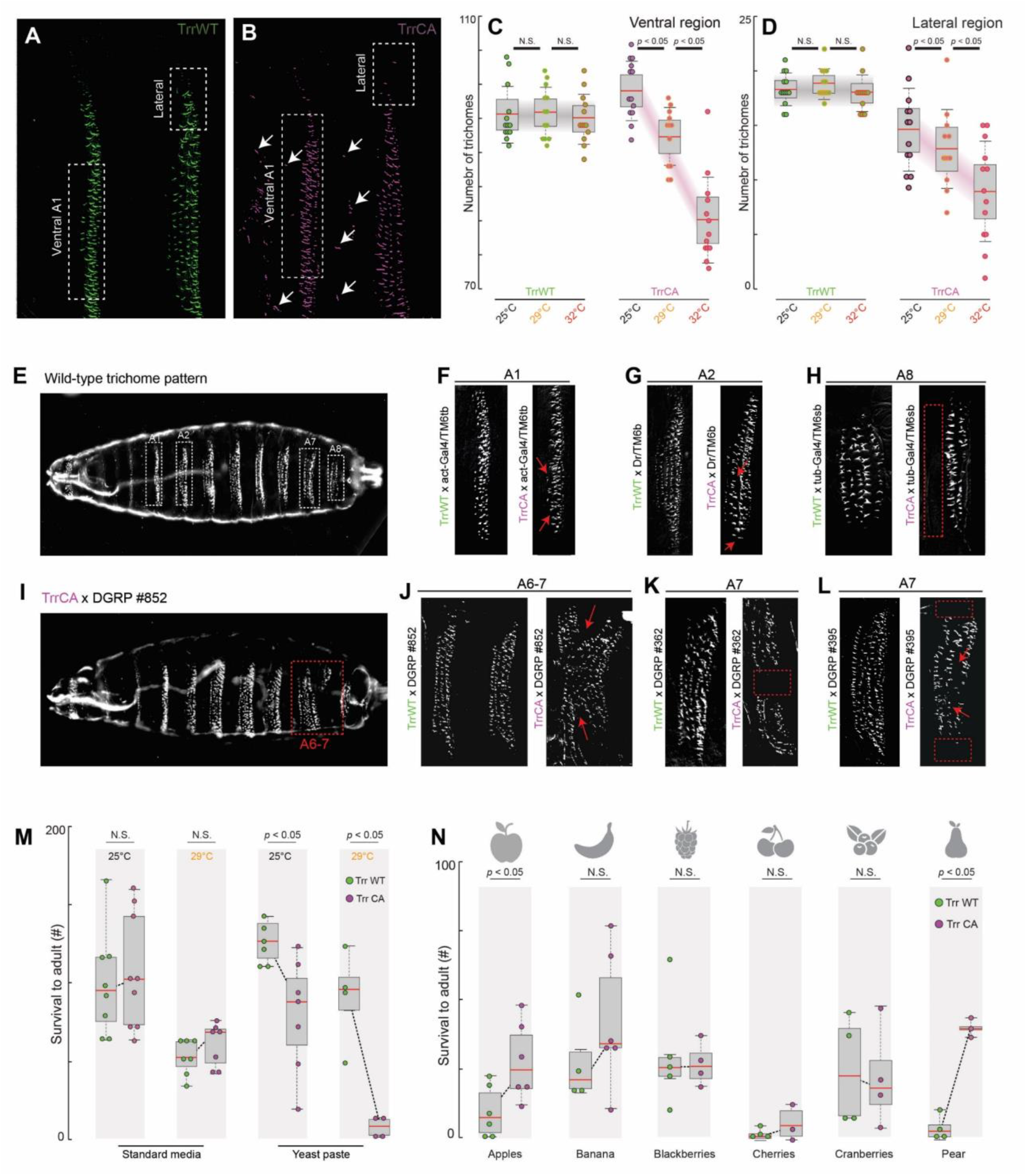
H3K4me1 conceals cryptic genetic variation and affects adaptability to specific environmental conditions. Related to Supplementary Figure 2. (A & B) Trichome patterns of the first two abdominal segments at 25 °C in both *trr*^*1*^ lines. The white arrows highlight the presence of ectopic trichomes. (C & D) Number of trichomes in the ventral box and the lateral box, respectively, in both *trr*^*1*^ lines and at different temperatures. Number of larvae quantified: 13 TrrWT and TrrCA at 25 °C, 13 TrrWT and 12 TrrCA at 29 °C, and 13 TrrWT and TrrCA at 32 °C. N.S.: not significant. (E) Wild-type trichome pattern as observed through dark field microscopy, with boxes highlighting the specific abdominal segments affected in the different crosses. (F-L) Details of specific abdominal segments (A1, A2, A6, A7 or A8) of cuticle preparations highlighting the trichome defects. The trr1 mutant lines were crossed with (F) act-gal4/TM6tb, (G) Dr/TM6b, and (h) tub-Gal4/TM6sb balancer stocks, or the (I & J) DGRP #852, (K) DGRP #362, or (L) DGRP #395 lines. (M) The number of offspring flies produced by both *trr*^*1*^ mutant lines with equal numbers of parental flies in two weeks, on standard lab medium (left) or lab food enriched with yeast paste (right), and at 25°C or 29°C. Each dot represents an independent replicate population. The boxed region is one s.d. and the tails are two s.d. (95 %). (N) Similar experiment to (M), but carried out in poorer food sources produced from slightly rotten organic fruits. The boxed region is one s.d. and the tails are two s.d. (95 %).

### H3K4me1 hypomethylation revealed hidden genetic variations

We then tested if H3K4me1 deposition maintains robust phenotypes by buffering not only against environmental stimuli but also different genetic variants. We outcrossed the TrrCA and TrrWT lines with different genetic backgrounds (see *Morphological analysis of larvae and adult flies* in **Material and Methods**), picking three balancer lines, whose lack of recombination could impair the purging of slightly deleterious genetic variants^24^. We analyzed the trichome patterns in larvae generated from these crosses. In all cases, we observed increased frequencies of aberrant trichome patterns with H3K4me1 hypomethylation (Figure 2E-H and Supplementary figure 2A). To increase the range of tested genotypes, we performed crosses with three lines from the Drosophila Genetic Reference Panel (DGRP), where the standing genetic variation represented in these lines could lead to additional altered phenotypes^25^. We found an increased frequency of altered trichome patterns produced by hypomethylation in these genetic backgrounds (Figure 2I-L and Supplementary figure 2A; p < 0.05, Chi-Squared goodness of fit test). The specific alterations, both in terms of the affected abdominal segment and the trichome rows that were missing/modified, were genotype-specific (Figure 2E-L).

Furthermore, TrrCA adults from some of these crosses showed increased wing defects, with one or both wings crumpled (Supplementary Figure 2B-E; p < 0.05, Chi-Squared goodness of fit test). Only crosses between TrrCA and specific genotypes showed higher penetrance of this aberrant wing phenotype compared to TrrWT (Supplementary Figure 2E), again suggesting a genotype-specific effect of hypomethylation. Thus, these results suggest that H3K4me1 can conceal genetic variation, leading to higher morphological homogeneity in a population, which would be consistent with the role of a phenotypic capacitor^24^.

### Hypomethylation altered adaptability to environmental perturbations

Phenotypic capacitors are proposed to facilitate the emergence of novel features, which are revealed under specific conditions of challenge to the organism^26^. To test this hypothesis, we analyzed if H3K4me1 hypomethylation can alter the adaptability to non-standard feeding regimes. We set up mating groups (20 females and 10 males) from the TrrCA or TrrWT lines in vials containing different food sources: standard lab food (molasses-based), standard lab food supplemented with yeast paste, or several media produced from organically grown fruits. The number of eclosed adults after two weeks was similar for TrrCA and TrrWT under standard feeding conditions, even at 29 °C (Figure 2M, left). In contrast, supplementing this food source with yeast paste led to a decrease in the offspring number of TrrCA compared to TrrWT. Increasing the temperature to 29 °C exacerbated this effect (Figure 2M, right). Non-standard fruit-based food sources yielded more heterogeneous results. Strikingly, in some cases the TrrCA line had increased offspring numbers. For example, the apple- and pear-based foods showed increased progeny (*p* = 0.027 and *p* = 0.026, respectively) (Figure 2N). Thus, these results show that H3K4me1 alters adaptability in a food-dependent manner, to the point of sometimes increasing adaptability outside of standard laboratory conditions.

### Hypo-monomethylation of H3K4 led to increased adult and larval size

The wide distribution of H3K4me1 across the genome^6^ connects it to many active or primed enhancers. This dense connectivity to regulatory networks of different functions may have pleiotropic influence on complex traits. Thus, to understand the full impact of H3K4me1 hypomethylation on a developmental system, we analyzed phenotypes in these *trr*^*1*^ mutant lines that are the result of interactions between multiple regulatory and signaling networks.

A pronounced effect is that TrrCA adult flies were larger than control ones (Supplementary Figure 3A-C, *p*=0.0103). This observation is consistent with a previous work showing that *trr* restricts growth in a cell-autonomous manner^27^. However, the effects of H3K4me1 hypomethylation on the size of different body features on the adult fly had not been measured before. We employed a morphometric approach^28^ and measured the length of three features that are commonly employed to identify different morphs or species^29,30^: wing intervein length, the tibia length, and head width. All three structures showed an increase in size in hypomethylated flies (Supplementary Figure 3D and F-H, *p*=0.008 for wings, *p*=0.012 for tibiae, and *p*=0.024 for heads). The difference in thorax size, head width, and the wing intervein length (but not the tibia length) increased with temperature to which the populations were exposed during development (Supplementary Figure 3E-H).

We noted that this effect on size was not restricted to adult flies: hypomethylated larvae were also larger than control larvae at the same stage of development (Figure 3A-C, *p* < 0.01). However, pupariation time was not affected by the difference in larval size (Supplementary figure 4A). A possible explanation is that the modulation of H3K4me1 levels may alter lipid metabolism, a known regulator of larval size^31^.

**Figure 3.**
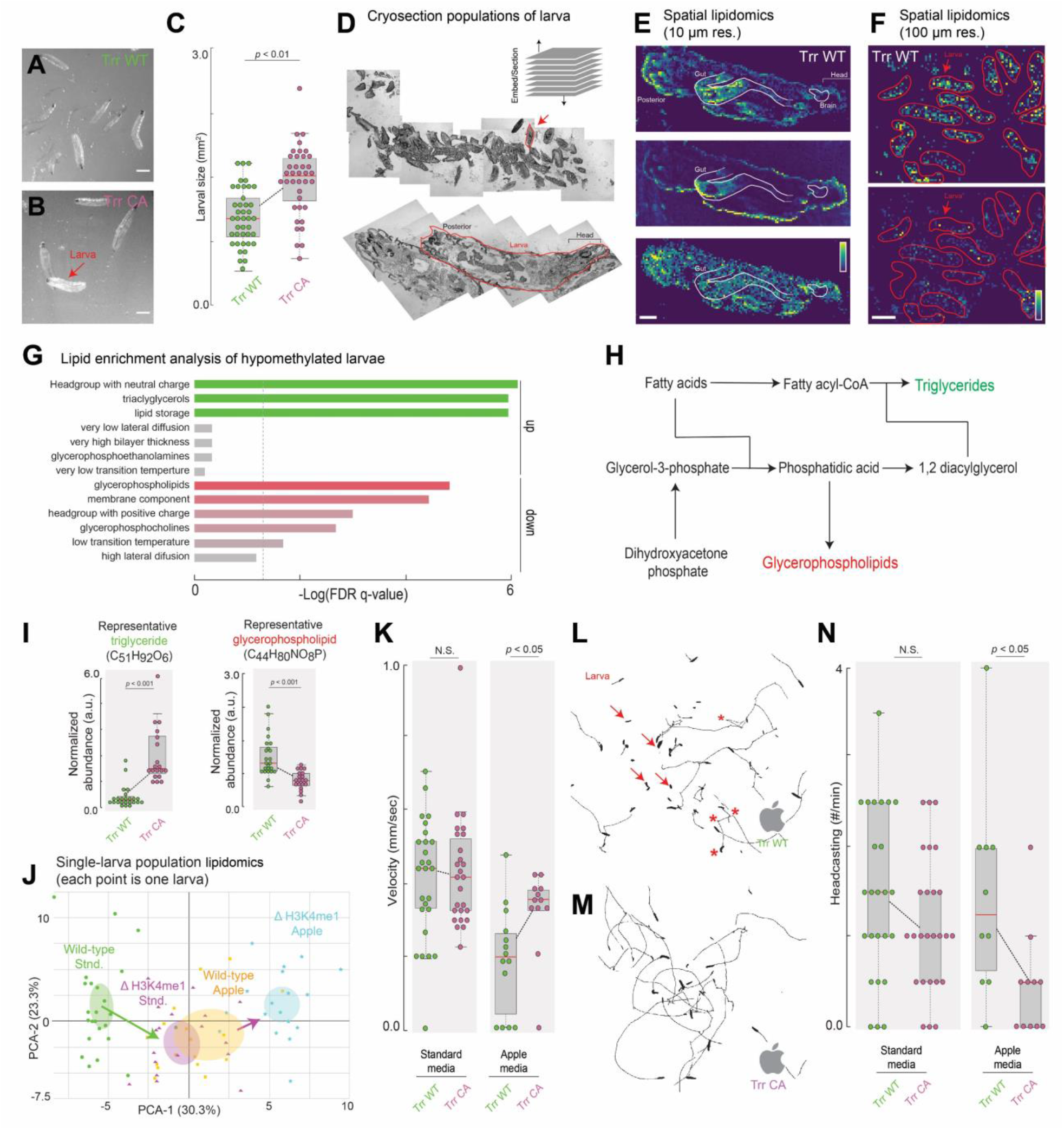
H3K4 hypomethylation alters metabolism and behavior. Related to Supplementary Figure 4. (A & B) Pictures of 120 h old larvae from TrrWT (A) or TrrCA (B) (scale bar = 830 µm). (C) The mean size of TrrCA larvae versus TrrWT (n=41 for TrrWT and n=39 for TrrCA). Center line, mean; upper and lower limits, s.d.; whiskers, 95% CIs. One-tailed t-test comparing the two Trr1 lines. (D) Upper panel: Middle section of a small larval population. The red arrow highlights an individual larva. Bottom panel: Middle section of a single larva at higher magnification. (E) High spatial resolution MALDI-imaging analysis of a 3rd instar larva (scale bar = 100 μm). The images show relative intensities of individual lipid species, each for an individual m/z value: upper panel = 544.3373 (C26H52NO7P); middle panel = 177.0158 (C7H6O4); lower panel = 744.5537 (C41H78NO8P). (F) Medium-resolution MALDI-imaging analysis for a larvae population, showing relative intensities for a glycerophospholipid (upper panel) at m/z=744.5537 (C41H78NO8P) and a triglyceride (lower panel) at m/z=815.6525 (C49H92O6); scale bar = 1 mm. (G) Enrichment analysis comparing both trr1 mutant lines (TrrWT vs TrrCA), based on the abundance values for 77 metabolites consistently detected in all tested conditions (the full list is in Supplementary Figure 4B). (H) Schematic of lipid metabolism with two products highlighted: triglycerides (green) and glycerophospholipids (red). (I) Abundance values of a representative triglyceride (left) and glycerophospholipid (right) obtained by MALDI-imaging mass spec with single-larva resolution. Each dot represents an individual larva with n=24 for TrrWT and n=20 for TrrCA. The boxed region is one s.d. and the tails are two s.d. (95 %). (J) Principal Components Analysis (PCA) based on single-larva abundance values for 77 different lipids identified by MALDI imaging mass spec. Each dot represents an individual larva. n=23 for TrrWT standard, n=20 for TrrCA standard, n=17 for TrrWT apples and n=20 for TrrCA apples. (K) Average velocity of individual larvae grown on either standard lab food or apple-based food. (L & M) Two-minute trajectories of TrrWT (L) or TrrCA (M) larvae grown in apple-based food. Red arrows point to larvae that remained still throughout the recording. Red stars show the change in larval paths associated with the head casting behavior. (N) Frequency of head casting of both trr1 mutant lines, on standard or apple-based food. These measurements only considered larvae that were moving actively. For all panels in the figure: Centre line, mean; upper and lower limits, s.d.; whiskers, 95% CIs. Two-tailed t-test comparing the two trr1 lines; NS not significant. n=26 for TrrWT and TrrCA on standard food, n=14 for TrrWT on apples, and n=12 for TrrCA on apples.

### Mass-spectrometry of hypomethylated larvae revealed increased triglycerides synthesis

To test if H3K4me1 hypomethylation alters lipid metabolism, we used MALDI-imaging mass spectrometry, a technique for spatial lipidomics that can detect various lipids with spatial resolution^32^. Larval populations of TrrCA and TrrWT were cryosectioned to 20 μm-thick sections (Figure 3D) and analyzed by MALDI-imaging with 10 μm pixel size. Figure 3E shows representative images, every image showing relative abundances of a particular lipid across a larva section, demonstrating the detection of lipids associated with larval anatomy.

To acquire population-level data through MALDI-imaging, we used a lower spatial resolution of 100 μm pixel size (Figure 3F). In larvae exposed to standard lab food, hypomethylation increased triglycerides levels, concomitant with a reduced abundance of glycerophospholipids (Figure 3G-I, Supplementary Figure 4B). The elevated triglycerides concentration was confirmed by a biochemical assay (Supplementary Figure 4C).

We next tested the effect of hypomethylation when larvae were raised with an apple-based medium as a non-optimum, carbohydrate-rich food source, where hypomethylation increased adaptability (Figure 2N). A principal component analysis (PCA) of single-larva lipidomic profiles integrating intensities of 77 lipids detected across all conditions revealed that the effects of H3K4me1 hypomethylation is dependent on the feeding regime (i.e. Apple vs. Standard, Figure 3J, Supplementary figure 4B, D & E). In contrast to standard lab food, enrichment analysis showed that hypomethylated larvae raised on apples had increased levels of glycerophosphoethanolamines, with unaltered triglyceride abundance (Supplementary figure 4D). This population-level analysis of lipid signatures suggests that H3K4me1 hypomethylation widely alters larval lipid metabolism in a food-dependent manner.

### Hypomethylation altered larval behavior on non-standard food sources

Metabolic states can alter behavioral patterns in *Drosophila*^33^. Therefore, we measured the crawling velocity of larvae from both TrrCA and TrrWT that developed either on standard lab food or apple-based medium. We did not find differences in the mean speed between these two genotypes on standard lab food (Figure 3K left). However, TrrCA larvae on apple-based medium crawled at a higher velocity than TrrWT larvae (Figure 3K right, *p* = 0.014). Moreover, H3K4me1 hypomethylation altered the movement patterns of larvae, as evidenced by their crawling trajectories (Figure 3L & M). Thus, we also quantified the frequency of exploratory head casting, a stereotyped larval behavior^34^. Similar to crawling velocities, we only found differences on apple-based food (Figure 3N, *p* = 0.029). To verify that the effects that we observed in TrrCA are linked to hypomethylation, we generated a new fly line from *w*^*1118*^ using CRISPR/Cas9 to modify the native *trr* locus. We found that this CRISPR.TrrCA line (Supplementary Figure 5A-F) showed many of the phenotypes that we detected in the trr^1^;;TrrCA line (Supplementary Figure 5G-L), including similar behavioral alterations. This suggests that the effects of hypomethylation described here are consistent between populations and possible genetic background variations. In summary, reduction in H3K4me1 led to changes in larval behavior on food sources not commonly utilized in the laboratory but available in nature.

## Discussion

H3K4me1 is a canonical histone modification marking transcriptional enhancers across a wide array of genomes^35^. Despite its ubiquity, previous works have demonstrated that H3K4me1 deficiency is tolerated^12,17^ and that gene expression is mostly unaffected^17^. A possible explanation has been that it is a “fine-tuner” of enhancer functions—permitting more nuanced gene expression in response to environmental perturbations^17^. Furthermore, it has been proposed that chromatin regulators may have the ability to buffer gene expression variations, which might be a general characteristic of large-scale chromatin regulators^36^.

Here, we have approached its biological role through a phenomics approach^5^ across animal development—acquiring phenotypic data that range from gene expression to behavior. We have shown that H3K4me1 provides regulatory robustness to variable environments and genetic variations (Figure 4). For individual regulatory networks, it preserves correct gene expression and cell fate determination in the face of environmental stresses by supporting the enrichment of transcription factors in transcriptional microenvironments.

**Figure 4:**
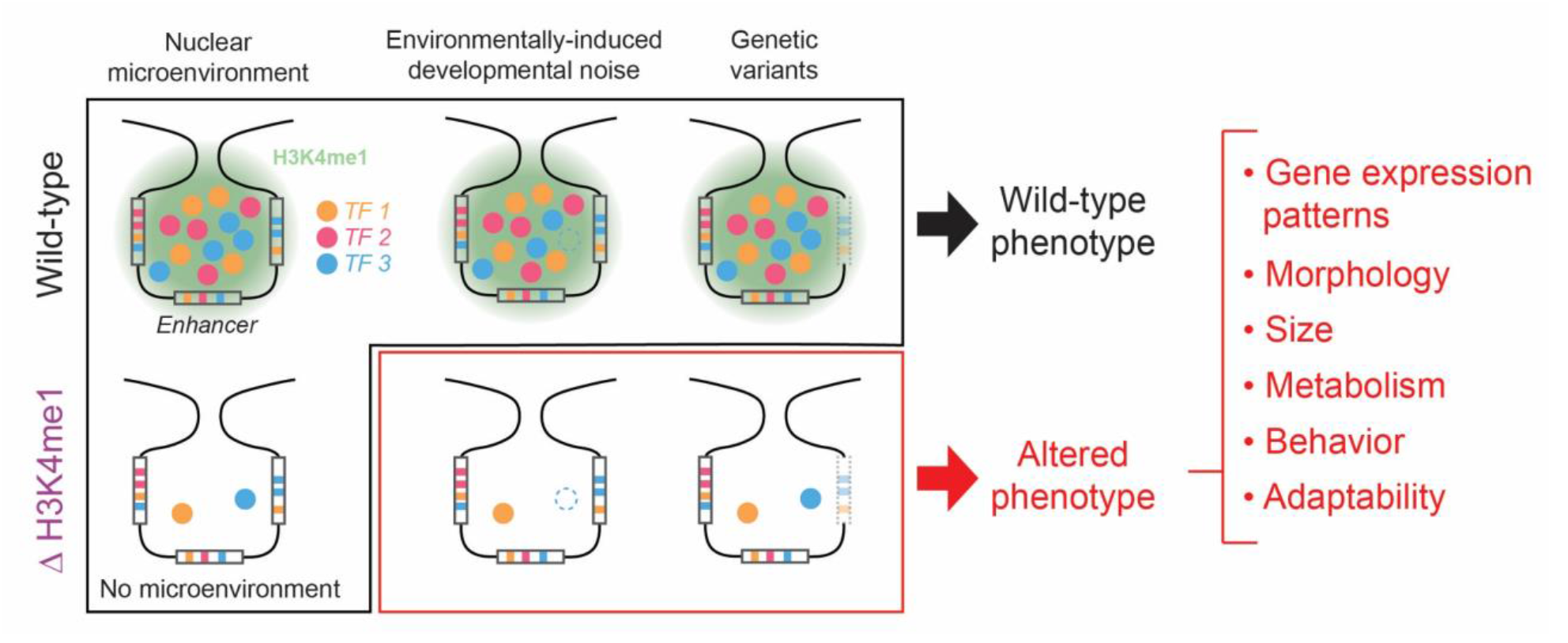
H3K4me1 stabilizes gene transcription through the establishment of nuclear microenvironments, allowing the emergence of alternative states. H3K4me1 may contribute to the establishment or maintenance of localized transcriptional environments. These dynamic structures facilitate stable transcriptional outputs, as the local clustering of TFs and enhancers can minimize the effect of environmental and genetic perturbations. The absence of H3K4me1 alters phenotypes at many different levels, leading to specific alterations in size, morphology, metabolism, behavior, and adaptability in context-dependent ways.

At a population level, H3K4me1 may conceal genetic variations that would otherwise cause unfavorable phenotypes, potentially functioning as a phenotypic capacitor^24^ (Figure 4). Importantly, H3K4me1 hypomethylation did not completely disrupt any analyzed phenotypes, but instead altered them in specific ways. For example, trichome and wing defects only appeared in specific genetic backgrounds (Figure 2E-L). Hypomethylation even increased adaptability to specific food sources (Figure 2N). The extensive range of phenotypic variation specific to inbreed lines and environments establishes that H3K4me1 has global effects on the storage of cryptic polymorphisms and their release in response to shifting environments. Together, these results support the role of H3K4me1 as a phenotypic capacitor to buffer intraspecies genetic variation, potentially linking this epigenetic mark with the emergence of novel traits.

The use of the developmental phenomics workflow introduced here allowed us to describe how impaired transcriptional robustness propagates across the entire biological system, altering every analyzed phenotype. Thus, this focus on phenotypic changes across developmental scales provided a mechanistic link between H3K4me1 and its role in fostering developmental robustness through transcriptional microenvironments (Figure 4). Even though they appear to be a central feature of gene expression, the regulatory mechanisms underlying these microenvironments and their resulting physiological implications are just starting to be explored^23^. DNA accessibility is usually considered a key element in the clustering of transcription factors and polymerases in transcriptional hubs^37^; however, evidence for a histone mark playing a role in the organization or maintenance of nuclear microenvironments had not been reported before. Future research could reveal the full extent of this phenomenon, as well as additional elements that might collaborate with H3K4me1 in the establishment of nuclear microenvironments.

The diverse effects of H3K4me1 can be conceptualized using the molecular framework shown in Figure 4. The biochemical function of H3K4me1, including the associated histone mark H3K4me3^38^, may be to guide multivalent proteins into transcriptional condensates or “hubs”^23^. In so doing, such histone marks may stabilize localized proteins concentrations and activate them in the proper place and time during development. Therefore, as originally proposed for Hsp90^24^, such marks may contribute to phenotypic variance by buffering the functional state of transcriptional enhancers—in this case, bound transcription factors, other histone marks, and local co-activator concentrations—that contribute to altered traits through the effects of chance, genotypes and environments^39^.

A novel component of this research was the use of MALDI-imaging mass spectrometry for the measurement of single-larva and population lipid profiles. Thanks to this analysis, we were able to reveal distinct metabolic profiles of larvae that outwardly appeared to be wild-type. The same approach can detect small molecules, small peptides, glycans, and exogenous molecules such as drugs or pollutants^40^. Thus, this opens the avenue towards fast and cost-efficient metabolic phenotyping at a population scale. Our general approach, combined with advances in robotics^41,42^ and automated behavioral characterization^43^, could turn phenomics into a standardizable phenotyping method for multiple fields of biological research.

In conclusion, this work highlights the risks of stripping away too much of how variable ecologies affects the function of animal genomes. A common thread that emerged from our investigations is that “standard laboratory conditions” turned out to be ill-suited for teasing out the widespread effects of H3K4 monomethylation. These results highlight that incorporating realistic ecological and environmental contexts into our experimental design is essential for understanding the regulatory genome and its contribution to evolution and development^44^. In the future, insights from phenomics and the inclusion of ecologically relevant conditions will allow us to explore how modulating elements embedded in densely connected biological networks could lead to the emergence of novel traits and influence the evolutionary dynamics of entire populations^45–47^.

## Acknowledgments

Albert Tsai is supported by the German Research Foundation (Deutsche Forschungsgemeinschaft, TS 428/1-1).

Mans Ekelöf and Theodore Alexandrov are supported by the ERC Consolidator grant METACELL. (grant agreement 773089).

Rafael Galupa and Lautaro Gandara are supported by fellowships from the European Molecular Biology Laboratory Interdisciplinary Postdoc Programme (EIPOD) under Marie Skłodowska-Curie Actions COFUND (Grant agreement numbers 664726 and 847543, respectively).

Albert Tsai, Mans Ekelöf, Theodore Alexandrov, and Justin Crocker are supported by EMBL. Ella Preger-Ben Noon is supported by the Israel Science Foundation (grant No. 2567/20).

The transgenic fly lines containing Trr methylation mutants were a generous gift from A. Shilatifard.

## Data and material availability statement

The original images (cuticle preparations and embryo images, organized into zip files) will be available for download and are indexed at: https://www.embl.de/download/crocker/XXXXX. All fly lines will be available upon reasonable request. Spatial lipidomics data is available through the METASPACE platform:

## Materials and Methods

### Fly strains and crosses

All fly strains were kept at standard laboratory conditions at room temperature unless otherwise noted. We used *w*^*1118*^ as the “wild-type” reference in the experiments shown in Figure 1A & B, Supplementary Figure 1B-E, and Supplementary Figure 5. Otherwise, we used lines with non-functional Trithorax-related allele (*trr*^*1*^) with two different Trr rescue constructs on the third chromosome: the wild-type rescue line (*trr*^*1*^*;;trr*(WT)) or the hypomethylated line (*trr*^*1*^*;;trr*(C2398A)). These lines were established and characterized in a previous work^17^.

For experiments examining larval and adult phenotypes with different genetic backgrounds, we crossed the *trr*^*1*^ lines with balancer stocks obtained from the Bloomington Stock center (https://bdsc.indiana.edu/index.html). They are: *;;Dr/TM6b (BS00211),; iso tub-Gal4 (VII)/TM6sb* (from Maria Leptin) and *;;act-Gal4/TM6tb (3954)*. We also employed lines #362, #395 and #852 from the *Drosophila Genetic Reference Panel* (http://dgrp2.gnets.ncsu.edu/).

### H3K4me1 ChIP-Seq

Stage 15 embryos from a line containing *E10::GFP* and *7::dsRed* transgenes were cross-linked, dissociated and isolated nuclei were immunostained with anti-GFP and anti-dsRed antibodies. Following staining with appropriate secondary antibodies, the E10::GFP and 7::dsRed nuclei, which constitute only 1.6% and 2.1% of the total input material, respectively, were isolated by fluorescence activated cell sorter (FACS, Supplementary figure 1B). Chromatin from 250,000 nuclei of each cell sub-populations was isolated and used for ChIP with anti-H3K4me1 and anti-H3 antibodies (abcam) using the iDeal ChIP-seq kit from Diagenode. Libraries were prepared using the Ovation Ultralow V2 DNA-Seq library preparation kit (NuGen) according to the manufacturer instructions. Following sequencing adapters and low-quality reads (< Q20) were trimmed using TrimGalore (http://www.bioinformatics.babraham.ac.uk/projects/trim_galore). Mapping was performed with bowtie2 ^48^ using the reference genome dm6 and sensitive end-to-end presets. Unmapped, multi-mapping reads, reads mapping to chrM (and other non-standard chromosomes) and duplicate reads were removed. For normalization, we subtracted bigWig files of H3 ChIP-seq samples from bigWig files of H3K4me1 ChIP-seq samples. For visualization purposes, we averaged normalized replicates (Pearson correlations of 0.86-0.98) and normalized data was smoothened using a moving average smooth of 500bp.

### Correlation between H3K4me1 deposition and the regulation of the *svb* network

Segmentation genes with significant H3K4me1 ChIP-seq peaks within 10 kb of the transcription start sites were identified using the modENCODE dataset H3K4me1; Embryos 12-16 hours embryonic data^35^ (ID 780).

### Sample preparation and staining for confocal imaging

Embryos for imaging were collected, fixed in 5% PFA and stained according to previous protocols^49^. To detect *svb* transcription, antisense RNA probes with DIG were made using the primers from Tsai *et al*., 2019^21^. For the staining of *svb* and H3K4me1, the samples were first stained for the histone modification following the IF protocol^22^, re-fixed in 5% PFA in PBT (PBS with 0.1% Tween 20) for 20 minutes, and then stained for *svb* following the FISH protocol. For all other experiments with *svb*, we followed the FISH protocol^22^.

The following primary and secondary antibodies were used (with dilution ratio in parentheses followed by the manufacturer and catalog number):

Primary antibodies

Rabbit anti-H3K4me1 (1:250): Merck, 07-436

Mouse anti-Ubx (1:20): Developmental Studies Hybridoma Bank, FP3.38-C

Sheep anti-DIG: (1:250) Roche, 11333089001

Secondary antibodies

Donkey anti-mouse Alexa 555 (1:500): ThermoFisher, A31570

Donkey anti-rabbit Alexa 488 (1:500): ThermoFisher, A21206

Donkey anti-rabbit Alexa 555 (1:500): ThermoFisher, A31572

Donkey anti-sheep Alexa 488 (1:500): ThermoFisher, A11015

Donkey anti-sheep Alexa 633 (1:500): ThermoFisher, A21100

Stained embryo samples were mounted in ProLong Gold + DAPI Mounting Media (Molecular Probes, Eugene, OR) on a glass slide covered with a number 1.5 high precision coverslip.

### Confocal image acquisition and analysis

Confocal images were acquired on a Zeiss LSM 880 confocal microscope (Zeiss, Germany) under a Zeiss Plan-Apochromat 63x/1.40 NA objective with the appropriate laser lines (405, 488 and/or 561 nm) using the Zeiss-recommended optimal resolution. Imaging processing to locate transcription sites and extract spatial data was performed in Fiji/ImageJ^50^ with native functions and the 3D ImageJ Suite plugin^51^. Subsequent data analysis was performed in MatLab (MathWorks, Natick, MA) to extract transcription site intensity and radial distributions^21,22^.

### Sample preparation to analyze larval phenotypes (cuticle preps etc.)

Cuticle preps were imaged on a phase-contrast microscope (Zeiss, Germany). The number of trichomes in the A1 ventral band between two sensory cells was counted using a find maximum function in Fiji and reported as “Ventral”, as previously described^21^. The number of trichomes in the lateral extremity of the ventral band where the *svb* enhancer *DG3* provides exclusive coverage was also counted and reported as “Lateral”, as previously described^21^.

### Morphological analysis of larvae and adult flies

Female virgins from the Trr1 mutant lines were crossed with males from different DGRP stocks or balancer lines (see *Fly strains and crosses*). To analyze the larval trichome pattern, these crosses were housed in egg collection chambers. Embryos were then collected from plates and placed in water, on which they developed at 29^◦^C overnight. Afterward, 1st instar larvae were treated according to standard protocols^52^ to prepare cuticles for analysis. Cuticle preparations were imaged using dark-field and phase-contrast microscopy (Zeiss, Germany). For the morphological analysis of adult flies, these crosses were placed on fresh vials at 29^◦^C. After 16 h of egg laying, adults were removed, and the egg-containing vials were left at 29^◦^C for 10 days. Then, the emerged male adults were anesthetized with CO_2_, and wings were analyzed and photographed employing an Olympus stereoscope.

In all crosses, we used the *trr*^*1*^ mutant lines as the female parental strain, to make sure that all the males in the offspring were deprived of a wild-type Trr allele in the endogenous locus. As the female offspring is heterozygous for Trr1, and thus it is supposed to have normal H3K4me1 levels, it should be noted that the informed frequencies are likely an underestimation, and should be considered strictly in a qualitative manner.

### Offspring production assay with different temperatures and food sources

Populations of 2-day-old flies from the Trr1 mutant lines, consisting exactly of 20 females and 10 males, were placed in vials containing standard lab food, standard food supplemented with yeast paste, or food produced from slightly rotten fruits collected in a local forest (Heidelberg, Germany, GPS Coordinates, 49.38475495291698, 8.71066590019372). After 2 days of egg laying, adults were removed and the vials were placed at 25 °C. For lab food and lab food with yeast, replicates were carried out at 29 °C. Then, adult offspring were counted in each vial after 14 days.

As standard food, we employed a modified version of the BDSC Cornmeal Food (https://bdsc.indiana.edu/information/recipes/bloomfood.html), consisting of agar 40 g/l, dry yeast 18 g/l, soya powder 10 g/l, corn syrup 22 g/l, malt extract 80 g/l, corn powder 80 g/l, propionic acid 6.25 g/l and Nipagin 2.4 g/l. All fruit-based foods were prepared according to Chhabra *et al*., 2013 ^53^. Briefly, the indicated fruits were homogenized in a blender, and then water was added to a final concentration for the fruit mass of 1,5 g/l. After adding agar (10% m/v), these preparations were heated in a microwave oven and then dispensed into individual vials.

### Genotyping of the Trr locus in the 3^rd^ chromosome (rescue constructs) in mixed populations

To identify the Trr allele present in individual flies from the mixed populations, we used the following primers: TrrWT_fw (AGTCGCACAAGATACCGTGC), TrrWT_rv (TGCAATACAGTGGCAACGTC), TrrCA_fw (GCACAAGATACCGGCCG) and TrrCA_rv (CACGATACACGCAGCGAAAG). Both primer sets were used on each individual gDNA sample, so flies for which positive results were obtained with both sets were considered as heterozygous, whilst a single positive was identified as a homozygous fly. For each of the populations kept in standard food, 50 flies were genotyped per generation. In the case of the populations kept on apples, all living flies were genotyped in each generation.

### Larval lipidomics assays with MALDI-imaging

Larval tissues were cryo-sectioned before subjecting them to MALDI imaging mass spectrometry. To do this, a small population (n≈10) of 3^rd^ instar larvae were embedded in a previously heated 5% m/v carboxymethylcellulose (Sigma) solution. After solidification, the obtained molds were sectioned in a Leica CM1950 cryostat at -20C, producing slices with a thickness of 20 μm. These slices were then mounted on regular glass slides, always aiming to preserve the middle section (40-60 μm) of the sectioned larvae.

Uniform coating of tissue sections with microcrystalline matrix material is essential for MALDI-MSI. To process the larval tissues, a 2,5-dihydroxybenzoic acid (DHB) matrix (Sigma Aldrich) 15mg/ml, dissolved in 70% acetonitrile, was applied onto the samples, mounted on regular glass slides, by using a TM-Sprayer robotic sprayer (HTX Technologies, Carrboro, NC, USA). Then, these glass slides containing the larval tissues were mounted onto a custom slide adaptor and loaded into the MS imaging ion source (AP-SMALDI5, TransMIT GmbH, Giessen, Germany). Generated ions were co-axially transferred to a high mass-resolution mass spectrometer (QExactive Plus mass spectrometer, ThermoFisher Scientific). Positive mode MS analysis was carried out in the full scan mode in the mass range of 200-1100 m/z (resolving power R=140000 at m/z 200). Metabolite annotation was performed using the METASPACE cloud software^54^. The Principal Component Analysis of these results was performed on R using the FactoMineR and factoextra packages (http://factominer.free.fr/). Abundance values were batch-corrected using the ComBat method^55^. Enrichment analysis were carried out using LION/web^56^.

### Triglycerides quantification assay

The concentration of triglycerides in *Drosophila* larvae was measured using the Triglyceride Quantification Colorimetric Kit from Sigma (Cat. # MAK266). Ten, 120 h old (3^rd^ instar), larvae from either the TrrWT or TrrCA line were homogenized in an Eppendorf tube on a Nonidet P40 Substitute (Sigma, Cat. # 74385) 5% solution. Then, the triglycerides concentration of each sample was quantified following the instructions provided by the manufacturer. Absorbance was measured at 570 nm. All metabolic determinations were carried out on larvae that came from vials with the same larval density (30 larvae per vial), to avoid effects of crowding on metabolism.

### Larval behavioral assays

Larvae from both *trr*^*1*^ mutant lines, either grown in standard lab food or apple-based food, were placed on agar plates, and their movement was recorded using a regular webcam (Logitech, 1080p, 30 Hz) for two minutes. Then, the speed of individual larvae was calculated from their displacement in the x- and y-axes, which was obtained using the MTrack2 tracking algorithm (ImageJ). The frequency of head casting for individual larvae was manually measured in each of these videos.

### New TrrCA allele developed with CRISPR/Cas9

We cloned two trr DNA sequences, one upstream and the other downstream to the catalytic domain, to act as RNA guides for CRISPR/Cas9 mediated transgenesis, into the pCFD4 plasmid using FSEI and BBSI. In parallel, we synthesized a trr DNA sequence that includes the above-mentioned guides, but altering the nucleotides that are required to replace a Cys by an Ala at position 2398, to act as template. Silent mutations were also added to prevent Cas9 for recognizing and cutting this new sequence. This new construct was cloned in the pUC57 plasmid. Both construct-containing plasmids were then injected into a fly line that expresses Cas9 exclusively in the female germ line (Bloomington #51324).

Putative transgenic flies were crossed with *w1118* ones, and sequenced. After multiple crosses with this *w1118* line to homogenize the genetic background, homozygous TrrCA;; lines were established.

The sequences of the three constructs can be found in Supplementary Table 1.

