## Supplementary Material for "Developmental phenomics suggests that H3K4 monomethylation catalyzed by Trr functions as a phenotypic capacitor"

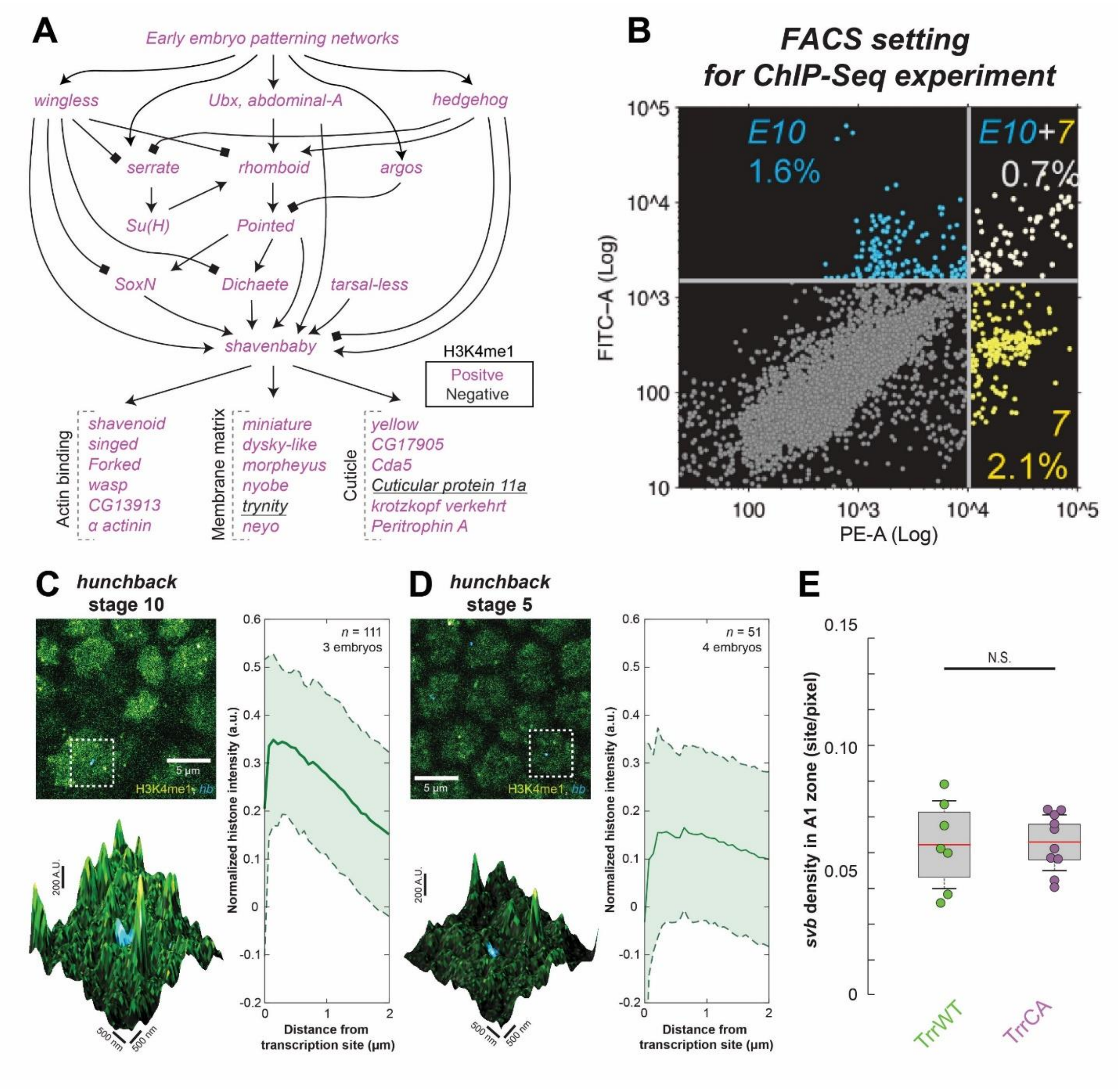


**Supplementary Figure 1. H3K4me1 and *svb* transcriptional robustness. Related to Figure 1.**

(A) H3K4me1 deposition near developmental transcription factors in the *shavenbaby* regulatory network (magenta indicates “positive” peaks within 10 kb of the start site).

(B) The setting used to sort cells in FACS for the ChIP-Seq experiment shown in Figure 1B.

(C-D) High resolution confocal imaging experiments in $w^{1118}$ embryos showing the distribution of H3K4me1 in *hb* transcription sites (C for stage 10 and D for stage 5 embryos). The lower panels show a zoomed-in view of the dotted boxes with the height indicating the intensity of the H3K4me1 signal. The normalized average H3K4me1 intensity over multiple transcription sites (C: n=111; D: n=51) is shown on the plots at the right. The shaded region is one standard deviation (s.d.). C & D were adapted from Tsai and Crocker, 2022 [20].

(E) Density of *svb* transcription sites in the first abdominal (A1) segment of stage 15 embryos at 25 °C within the ventral band where *svb* is normally expressed (compare with Figure 1H). The number of embryos are: 10 for TrrWT and 7 for TrrCA. The boxed region is one s.d. and the tails are two s.d. (95 %).

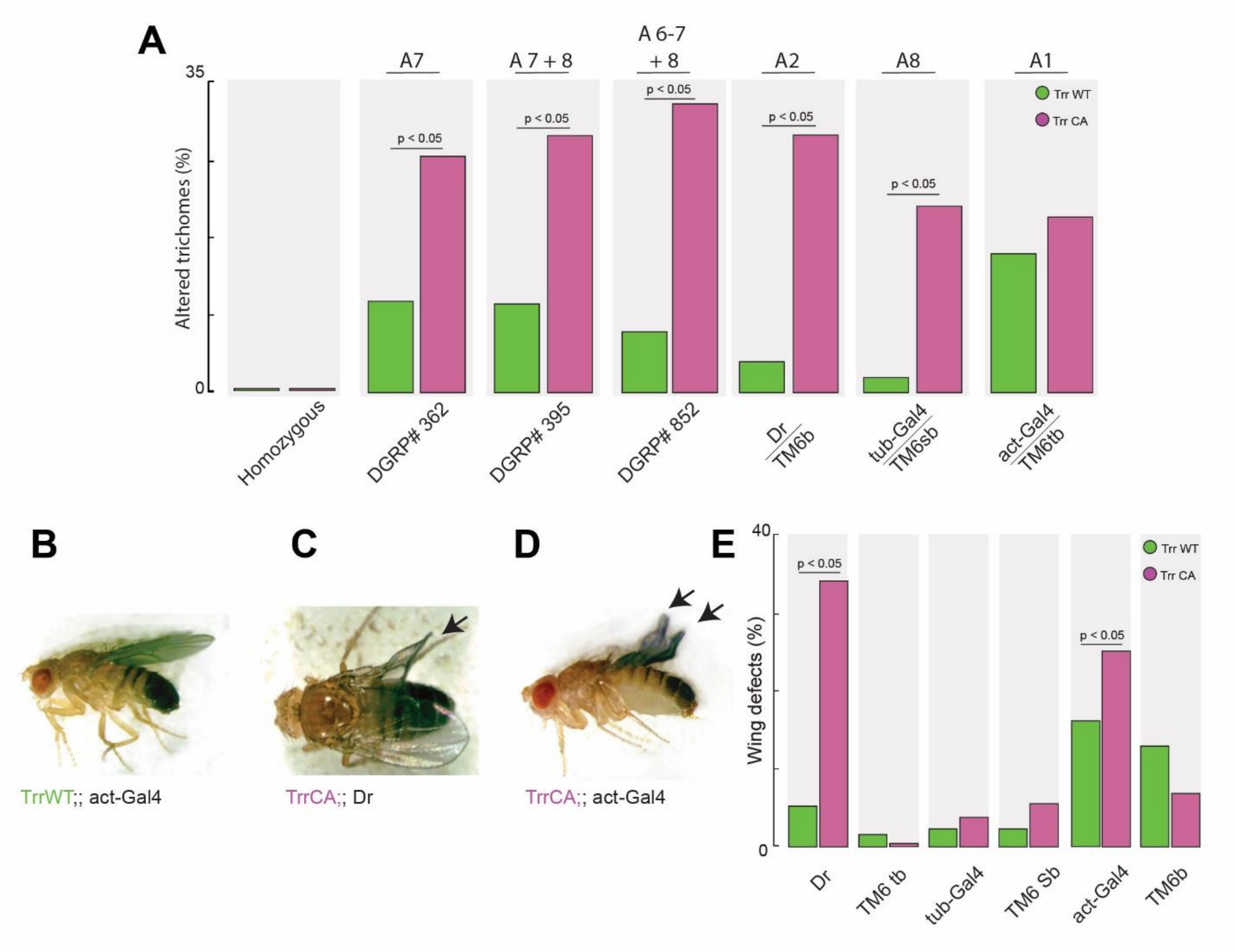


**Supplementary Figure 2: Aberrant morphologies in hypomethylated flies with different genetic backgrounds. Related to figure 2.**

(A) Frequencies of altered trichome patterns detected in the offspring of the crosses of the *trr*$^1$ mutant lines with the DGRP lines or balancer stocks. Only the genotype-specific alterations shown in Figure 2E-L were considered for the quantification. n=52 for TrrWT (homozygous), n=35 for TrrCA (homozygous), n=54 for TrrWT x DGRP#362, n=27 for TrrWT x DGRP#395, n=56 for TrrWT x DGRP#852, n=59 for TrrCA x DGRP#362, n=11 for TrrCA x DGRP#395, n=48 for TrrCA x DGRP#852, n=113 for TrrWT x TM6/act-G4, n=156 for TrrWT x TM6/tub-G4, n=45 for TrrWT x TM6/Dr, n=62 for TrrCA x TM6/act-G4, n=51 for TrrCA x TM6/tub-G4, n=57 for TrrCA x TM6/Dr.

(B-D) Pictures of adult *trr*$^{1}$ mutant flies with normal wings from TrrWT, or aberrant wing morphologies (arrows) from TrrCA.

(E) The fraction of flies with deformed wings observed in each of the previously mentioned crosses. n=116 for TrrWT;;act-gal4, n=112 for TrrWT;;TM6 tb, n=152 for TrrWT;;tub-Gal4, n=153 for TrrWT;;TM6 Sb, n=117 for TrrWT;;Dr, n=40 for TrrWT;;TM6b, n=34 for TrrCA;;act-gal4, n=72 for TrrCA;;TM6 tb, n=48 for TrrCA;;tub-Gal4, n=93 for TrrCA;;TM6 Sb, n=19 for TrrCA;;Dr, n=26 for TrrCA;;TM6b

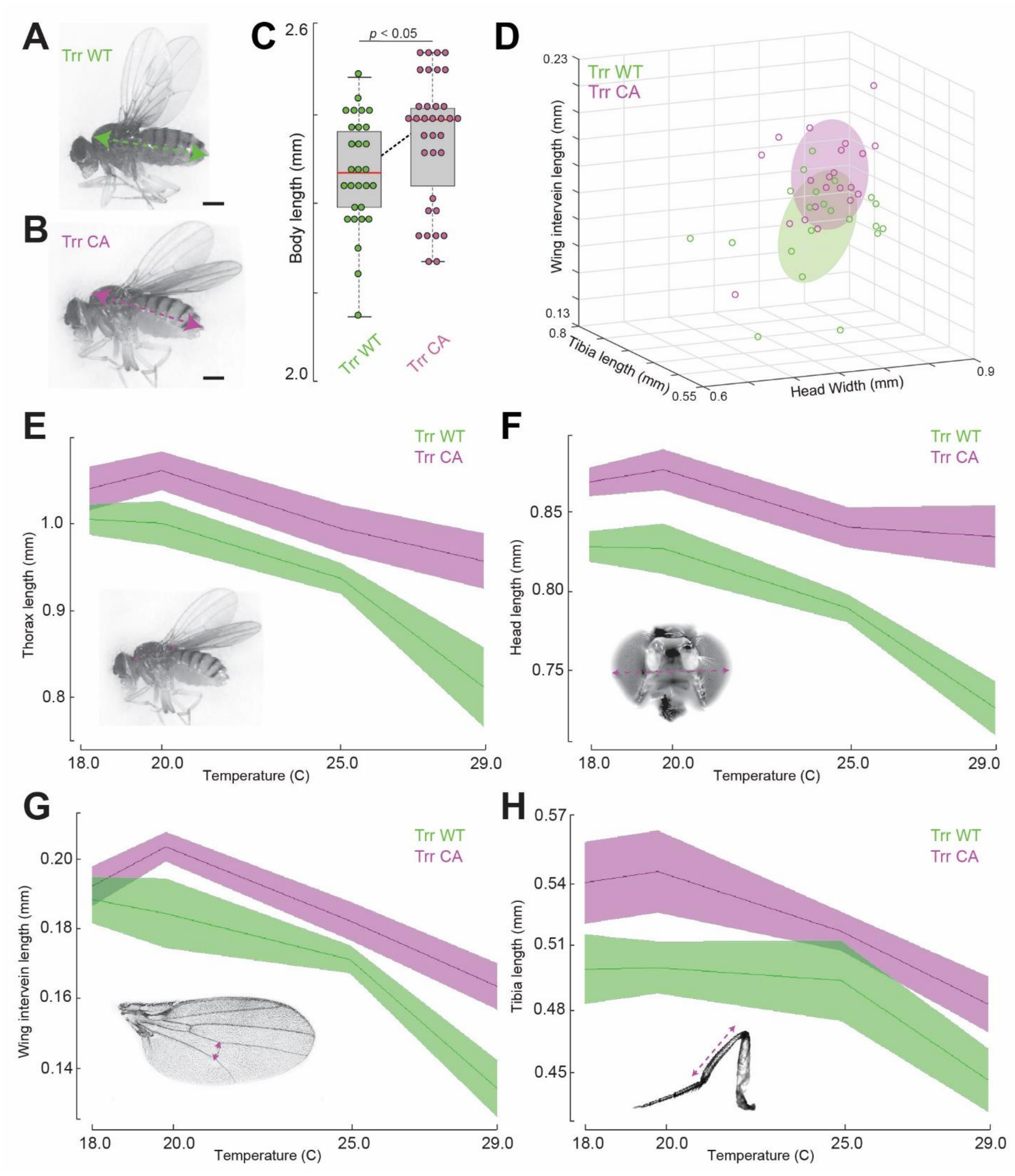


**Supplementary Figure 3: H3K4me1 controls body size in adults.**

(A & B) Pictures of 72 h old (after emergence) adult flies from TrrWT (A) or TrrCA (B) (scale bar = 0.5 mm). The dashed lines show the measured body length.

(C) The body length of TrrCA flies versus TrrWT (n=29 for TrrWT and n=34 for TrrCA). Center line, mean; upper and lower limits, s.d.; whiskers, 95% CIs. Two-tailed t-test comparing the two Trr1 lines.

(D) 3D plot showing the values from panels d to f but linked to single individuals from TrrWT or TrrCA (n=21). The ellipses show the 95% CIs for each population.

(E - H) Length of the thorax (E), the head width (F), the posterior cross-vein (G), and the fore tibia (H) in both TrrWT and TrrCA adult flies developed at different temperatures(n=20). Center line, mean; upper and lower limits, s.d.; whiskers, 95% CIs.

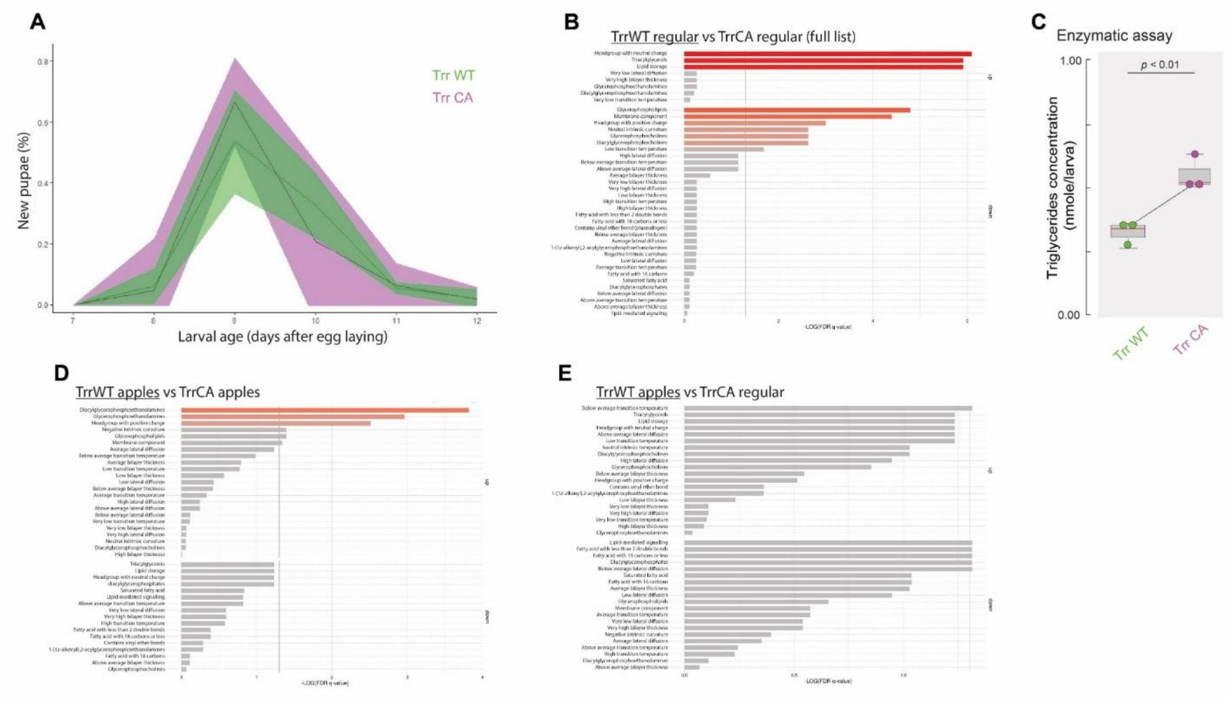


**Supplementary Figure 4: Metabolomics of the *trr*$^{1}$ lines. Related to Figure 3.**

(A) Pupariation time in both *trr*$^{1}$ mutant lines. The middle line shows the mean value for that time point, whilst the ribbon describes the CI (α=0.05); N (number of tested populations per condition) = 3.

(B) Enrichment analysis comparing both *trr*$^{1}$ mutant lines, based on the abundance values for 77 metabolites consistently detected in all tested conditions (full list)

(C) Enzymatic determination of triglyceride levels in TrrWT and TrrCA larvae with the same time of development (120 h). N=3.

(D & E) Enrichment analysis from pairwise comparisons between *trr*$^{1}$ mutant lines on apple-based food (D) and between the two populations with strong overlap in the PCA, TrrWT on apples and TrrCA on standard food (E).

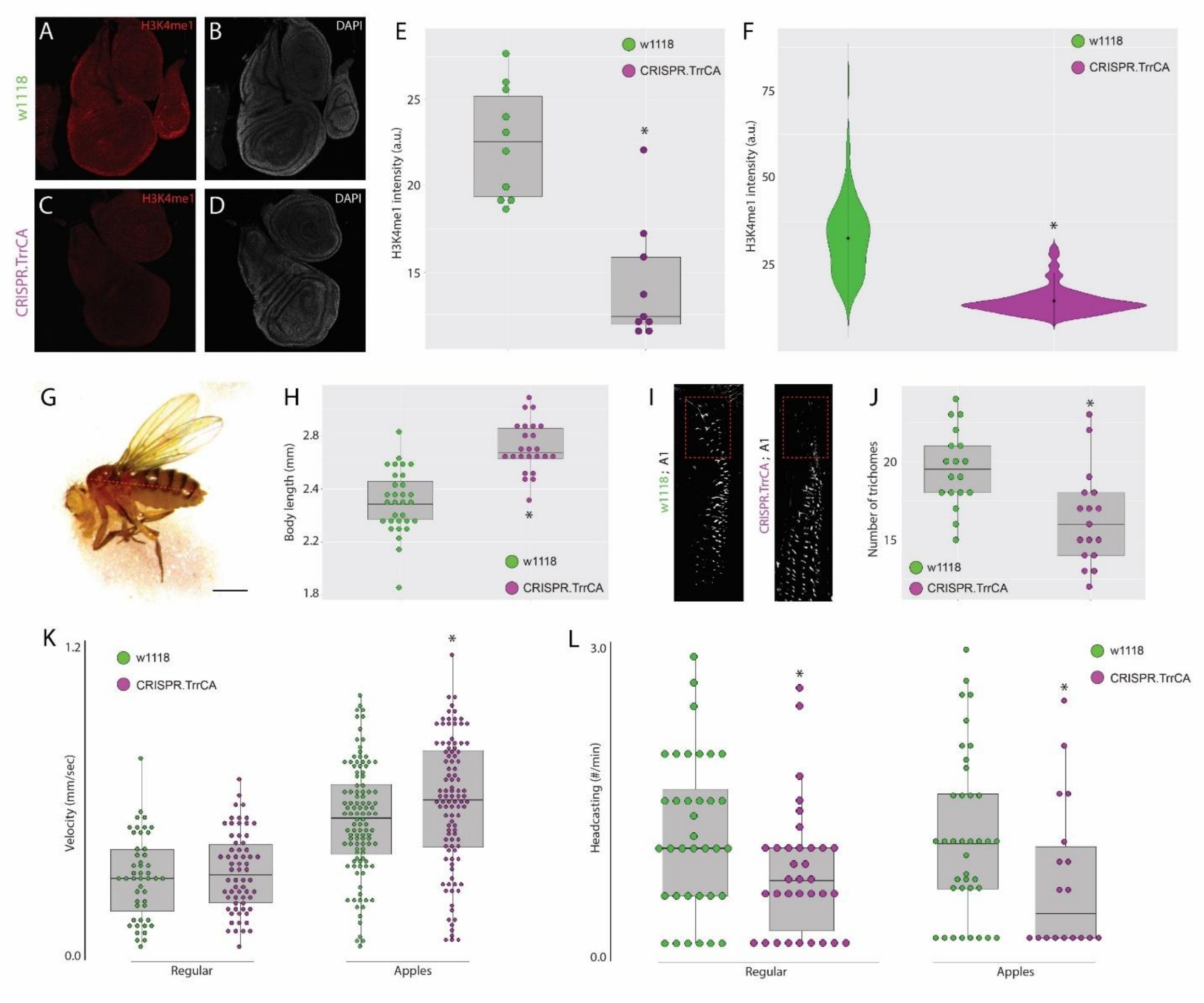


**Supplementary Figure 5. TrrCA mutation in the native locus (CRISPR.TrrCA) phenocopies TrrCA rescue construct.**

(A-D) Reduction in H3K4me1 levels in a CRISPR-built TrrCA allele (CRISPR.TrrCA) in the native locus. (A & C) Immunostaining showing the H3K4me1 signal, or (B & D) DAPI staining, in wing discs of w1118 or CRISPR.TrrCA.

(E-F) Intensity of the H3K4me1 signal in wing discs of w1118 or CRISPR.TrrCA. (E) Each dot represents the average value for an entire wing disc (n=10). Centre line, mean; upper and lower limits, s.d.; whiskers, 95% CIs. *$p<0.05$; Two-tailed t-test comparing the two genotypes (F) Single-cell H3K4me1 intensity measure in wing discs of the specified genotype (n=125). The red dot is the mean and the bar is one s.d. *$p<0.05$; Two-tailed t-test comparing the two genotypes.

(G) Pictures of 72 h old (after emergence) adult flies from CRISPR.TrrCA (scale bar = 0.5 mm). The dashed lines show the measured body length.

(H) The body length of CRISPR.TrrCA flies versus w1118 (n=31 for w1118 and n=24 for CRISPR.TrrCA). Center line, mean; upper and lower limits, s.d.; whiskers, 95% CIs. Two-tailed t-test comparing the two Trr1 lines.

(I) Trichome patterns of the first abdominal segment at 29 °C in w1118 or the CRISPR.TrrCA line. The dashed box highlights the sensitive lateral region in which the number of trichomes was counted.

(J)  Number of trichomes in the lateral box in w1118 and CRISPR.TrrCA at 29 °C. Number of larvae quantified: 18 for w1118 and 17 for CRISPR.TrrCA.

(K) Average velocity of individual larvae grown on either standard lab food or apple-based food.

(L) Frequency of head casting of both trr1 mutant lines, on standard or apple-based food. These measurements only considered larvae that were moving actively.

| | |
|---|---|
| gRNA1 | GTCAAAGTTACAAGAAAACATGG |
| gRNA2 | GCTATATGCAGATATGCTCGTGG |
| Template | TCAGAGCTAAGCAACCTGCTCCGTGTGGGCAACATGACGTTCCTGAACGTGGGTCAGTTACTGCCGCATCAGC<br>TGGAGGCCTTCCACACACCCCACTACATCTACCCCATTGGCTACAAGGTGGTAAGTACAAGTACCAGCCGGTC<br>GGCCGCCATGTTTATGAACAGCAATCTTAAATCCCGTTCCTCTTCCTCCCCCATAGAGTCGCTACTACTGGTGCG<br>TGCGACGACCCAATCGTAGGTGTCGATACATTTGCAGTATCGCAGAGGCCGGATGCAAGCCCGAGTTTCGCAT<br>CCAGGTGCAGGACGCCGGCGACAAGGAGCCGGAGCGCGAGTTTCGGGGCAGTTCGCCCTCGGCGGTCTGGC<br>AGCAGATTCTGCAGCCAATCACGCGACTGCGTAAGGTGCACAAGTGGTTGCAGCTCTTTCCCCAGCACATCAG<br>TGGTGAGGACCTATTTGGCCTAACGGAACCGGCCATTGTGCGAATCCTGGAAAGCCTCCCGGGCATTGAGAC<br>GCTCACTGACTACCGCTTCAAGTACGGTCGCAACCCGCTTCTTGAGTTCCCGCTAGCCATCAATCCGTCTGGAG<br>CGGCGCGTACGGAACCCAAGCAACGACAGCTGCTCGTCTGGAGGAAGCCGCATACGCAACGCACGGCCGGCA<br>GCTGCAGCACGCAGCGGATGGCCAACTCGGCTGCGATTGCCGGAGAAGTGGCCTGCCCGTACAGCAAGCAGT<br>TCGTGCATTCGAAGAGCTCCCAGTACAAAAAGATGAAGCAGGAGTGGCGCAACAACGTCTACTTGGCGAGGT<br>GAGACATTAGGTCCTAGTCAATAACCACTGATTACTTAAGCTCTCATTTTCTAGATCTAAGATCCAGGGTCTTG<br>GCTTGTATGCAGCTCGGGACATTGAAAAGCACACCATGATAATCGAGTACATTGGCGAGGTTATCCGCACCGA<br>GGTCTCCGAGATACGCGAGAAGCAGTACGAGTCCAAGGTGGGTTTCGCGACCATGCTTCCTTGTAACTTTGAC<br>TAATCTCTGATTCCCCAGAACCGTGGCATCTACATGTTCCGGCTGGATGAAGACCGCGTGGTGGACGCCACTCT<br>AAGCGGCGGGTTGGCGCGCTACATCAACCATTCGTGCAATCCCAACTGTGTTACCGAGATAGTGGAGGTCGAT<br>CGCGACGTGCGGATCATAATATTCGCCAAGCGGAAGATCTATCGCGGCGAGGAGGTGGGTCCCGCAAATAGA<br>AGTGTTTCGCCTGTACCTTAACTCACAATATCTCTTAAATTCTGTGCTTTAGCTCTCGTACGACTACAAGTTCGAC<br>ATTGAGGACGAGTCGCACAAGATACCG**GCC**GCCTGTGGAGCGCCCAACTGTCGCAAGTGGATGAACTAGAGT<br>CGGGAGGAGCCTCTAGCGGAGGACCAAAGGAAGCGCAAGAAGCGGCGGCGAGCACGTCGCTCCCAGTGAAC<br>AGGAACAGGAGTAGAGACAGGAACAGGAACAGGACTTGTATCTGATCAAGTAGCTAGATTCGCTAAGCATTA<br>TAGTTCGTGTACATTTATATATATATATATATATATATATATATATATTTATATATATATATAGTTTATATGCTATATGC<br>AGATACGCCCGTGGAGAGCATATGACCTTGTAAGTTTTGTATATGTGCCCACTGTAGCCAATTCGTATCCTTTA<br>TATCGCTCTTAAGTGCAAAAGTGGAGAGTGTCTTCATAGTTTAAATTGAATAATTGCTATCGTTTCTACATAGA<br>AAGAATATAAATGTTACTGCTAGGACACGACCACCCCTGTAATGTATTTCGGACCCCCTCTCCTCACACAGTCG<br>CTGGAGCACCTAATTTTATTTGTTTTAGCACCATCTCCACATTCATCACTACCACCATTGCAATTATCGCGAGGA<br>AACTGACCGCCCCGATGGCCAACCCAAAACTTGACTTTATGCGTGTGTGCCTTCCCCTCTGTTTACCATTATTAT<br>TATTATCATAATTGTAGCGGTTATCTTTCAATATTCGGCCACTTTACATGGGGAGCGATCCTTCTTTCGCTGCGT<br>GTATCGTGCGCAACCTTCTTTGAATATATACACTGATAAGGATATGAACACCTACTTCTATTTTATTTTATTTAAA<br>TAGTTTGCGATTACTTTTAAGTACATTAACTAACTCCGAACAACGTGTATTTTTCATCCCCTTAAATAGTAGCTA<br>AGTAAATTTTATGCCACCATTTTACGCACGTGATCTTTTATTCCAACTTATTGCAAACGGCAAACTGGAAATTAC<br>GAAATGGGGACGACAGATATTGCACCTAAGATCTTCGCTTGGCCAGCGCCAGAAGCCTCTGGAGCTGGACTG<br>CGGCAGCCAGGGTGAGTATGTCGAAGTTCTTCAGGTACCAGGAGATGTTGTACATGATGTGGATGCGCCAGT<br>TCTTGTCGAGGAACGCTTTGTAGAAGTGGCTCATCTGGTCCGCGGACACTTCGCTGGTTCCGCTTTCCTTGTGC<br>AGGCGGATGAAGTCTTCCTTTTCCTGTAAACGAACCAATTCGAAAGAGACGACATTAGCGGACTAGAAAGAGA<br>TGGCACAGTGGTCGTGGTTGTACCTCGTAGAAGCGCTTATTGTGTTTTGT |

**Supplementary table 1. Sequences employed to build a new TrrCA allele using CRISPR/Cas9. Related to Supplementary Figure 5.**

Sequences for the two guides and the template DNA that were required to edit the native *trr* locus using CRISPR/Cas9. The nucleotides in red are the ones that were altered to turn the Trr protein catalytically deficient. Guide-related sequences in the template are shown in green.
